# Lipid droplets as substrates for protein phase separation

**DOI:** 10.1101/2023.06.28.546804

**Authors:** Advika Kamatar, Jack P.K. Bravo, Feng Yuan, Liping Wang, Eileen M. Lafer, David W. Taylor, Jeanne C. Stachowiak, Sapun H. Parekh

**Author notes:** Correspondence should be sent to and.

## Abstract

Membrane-associated protein phase separation plays critical roles in cell biology, driving essential cellular phenomena from immune signaling to membrane traffic. Importantly, by restricting diffusion to a two-dimensional surface, lipid bilayers can nucleate phase separation at far lower concentrations compared to those required for phase separation in solution. How might other intracellular lipid substrates, such as lipid droplets, contribute to nucleation of phase separation? Distinct from bilayer membranes, lipid droplets consist of a phospholipid monolayer surrounding a core of neutral lipids, and they are energy storage organelles that protect cells from lipotoxicity and oxidative stress. Here, we show that intrinsically disordered proteins can undergo phase separation on the surface of synthetic and cell-derived lipid droplets. Specifically, we find that model disordered domains, FUS LC and LAF1-RGG, separate into protein-rich and protein-depleted phases on the surfaces of lipid droplets. Owing to the hydrophobic nature of interactions between FUS LC proteins, increasing ionic strength drives an increase in its phase separation on droplet surfaces. The opposite is true for LAF1-RGG, owing to the electrostatic nature of its interprotein interactions. In both cases, protein-rich phases on the surfaces of synthetic and cell-derived lipid droplets demonstrate molecular mobility indicative of a liquid-like state. Our results show that lipid droplets can nucleate protein condensates, suggesting that protein phase separation could be key in organizing biological processes involving lipid droplets.

## Introduction

Protein phase separation or condensation is the process by which proteins in solution self-assemble into a protein-dense phase by forming a network of weak, multivalent interactions (1). This process results in partitioning of proteins into a protein-rich phase and a protein-depleted phase. Under appropriate stimuli, including temperature, pH, ionic strength, and protein concentration, the protein-dense phase can behave like a liquid, such that droplets of this phase can coalesce, fuse, and undergo rapid molecular exchange with the protein-depleted phase (2). When the protein dense phase has the properties of a liquid, the assembly of condensates is referred to as liquid-liquid phase separation (LLPS). This process underlies the formation of membrane-less organelles like nucleoli, stress granules, and centrosomes, and is thus crucial for intracellular organization (3).

Recent work has shown that protein interaction with lipid membranes can induce LLPS at concentrations nearly an order of magnitude lower than would occur if proteins were free in solution (4–8). This difference in saturation concentration is important because the cytosol of the cell is densely populated by membrane-bound compartments, from organelles to trafficking vesicles (9). Therefore, lipid bilayers provide abundant substrates upon which LLPS could occur. For example, recent work has shown that formation of a liquid-like protein assembly at the plasma membrane is critical for productive endocytic events (6), and LLPS on giant unilamellar vesicles has been shown to drive membrane bending, suggesting that condensates could contribute to diverse membrane protrusions (4). Further, membrane-associated LLPS seems to be a broadly observed mechanism by which cells can organize complex processes at the membrane, including the force-dependent assembly of focal adhesions (10) and actin polymerization (7), synaptic vesicle clustering (11), synaptogenesis (12, 13), and T cell receptor signal transduction (14).

In addition to bilayer membranes, lipid droplets (LDs) are also ubiquitous lipid substrates found in the cytoplasm of most cell types. A lipid bilayer is composed of two leaflets of amphipathic lipids with the hydrophilic head groups oriented towards the aqueous environment and the hydrophobic tails pointed inwards toward each other. In contrast, an LD is composed of an amphipathic phospholipid monolayer surrounding a hydrophobic core composed of neutral lipids, including triacylglycerols (TAGs), sterol esters (SEs), and retinyl esters (15). LDs often have proteins embedded at the phospholipid surface, including the perilipin family (PLIN1-5), membrane trafficking proteins like Rabs, SNAREs, and Arfs, and proteins involved in protein degradation (16). While first understood as energy storage sites, in the past three decades, it has become clear that LDs are dynamic organelles with important roles in energy homeostasis, cellular communication, and disease (15). If LLPS helps to organize proteins on the surfaces of lipid bilayers, could it also occur on the surfaces of LDs? If so, it could provide a mechanism for organizing key events in LD biogenesis and function. Therefore, we investigated the extent to which LDs can provide a substrate for LLPS of proteins. Specifically, we characterized the condensation of well-studied proteins involved in LLPS on the surfaces of synthetic and cell-derived LDs. We found that at various salt concentrations, the low complexity domain of fused in sarcoma (FUS LC) and the RGG domain of LAF-1 (RGG), followed opposing trends in their tendency to condense on the LD surface, in line with the current understanding of the intermolecular interactions driving their condensation. Further, we found that these protein condensates remain liquid-like on the surface of both synthetic and cell-derived LDs. Our findings indicate that LLPS is a viable mechanism for protein organization on the LD surface.

## Results

### Formation and characterization of synthetic lipid droplets

Synthetic lipid droplets were produced according to previously published protocols (17, 18), with several modifications. Briefly, phospholipids were added to neutral triacylglycerols, and the solvent was evaporated by drying with N_2_. The phospholipid-oil mixture was heated to 37°C and vortexed to resuspend dried phospholipids in the oil phase. Then, an aqueous buffer was added, and the solution was vortexed to create an emulsion of phospholipid-coated lipid droplets in the aqueous buffer (Fig. 1A, B). To confirm that the resulting lipid substrates had the same structure as a cellular LD, we doped the phospholipids with 0.1 mol% Texas Red-DHPE and added Bodipy 505/515, a neutral lipid dye, and observed the respective localization of the dyed substrates using confocal fluorescence microscopy. As expected, in individual confocal slices, we observed a phospholipid ring surrounding a neutral lipid core (Fig. 1C, upper panel) in contrast to vesicles with a phospholipid bilayer surrounding an aqueous core (Fig. 1C, lower panel). To further investigate whether the phospholipid layer was truly a monolayer, as is characteristic of cellular LDs, we performed cryo-electron microscopy (cryo-EM) to visualize the LD structure at the molecular level. Electron micrographs revealed that the LDs were composed of a monolayer of phospholipids (Fig. 1D, upper panel) in contrast to vesicles that show two visible layers with a total thickness of approximately 5 nm, in good agreement with the thickness of DOPC vesicles previously reported (19, 20) (Fig 1D lower panel, Fig. S1).

**Figure 1:**
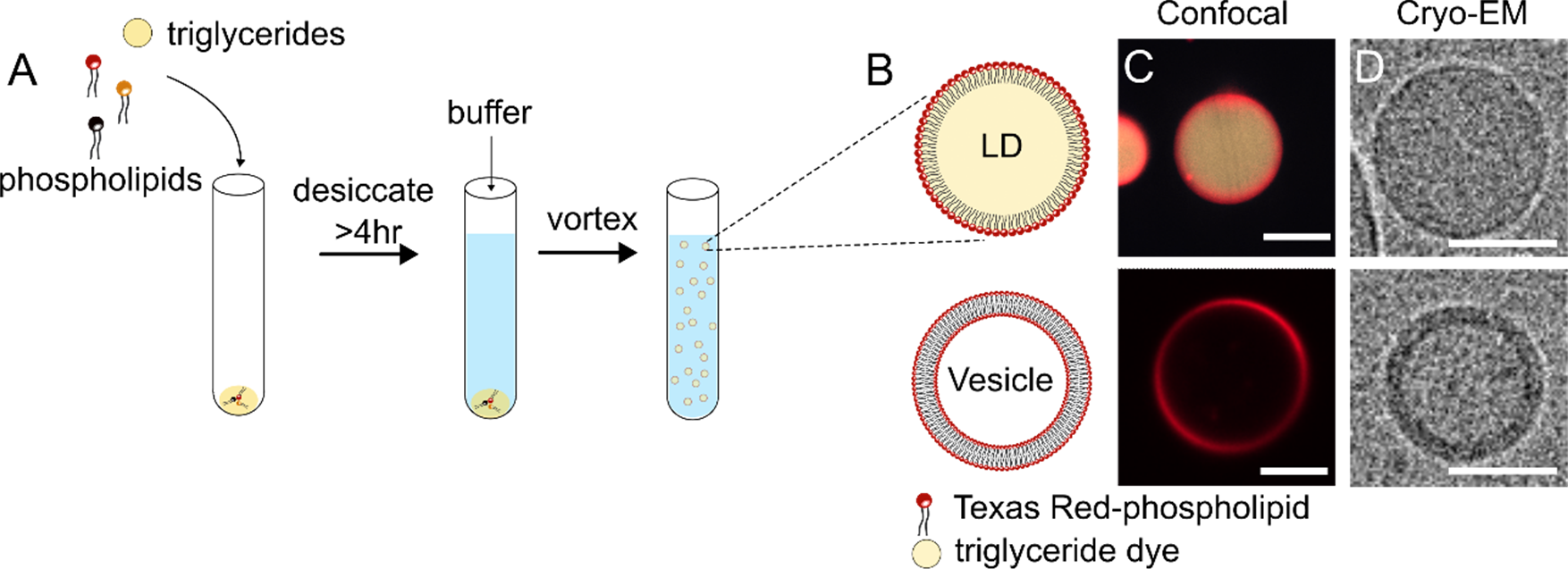
Lipid droplet (LD) synthesis and characterization. (A) Schematic of LD synthesis. (B) Schematic of an LD characterized by a monolayer of phospholipids surrounding a core of neutral triacylglycerol (TAG) (upper), and a bilayer vesicle characterized by a phospholipid bilayer surrounding an aqueous core (lower). (C) Confocal images of a representative synthesized LD composed of 98 mol% DOPG, 2 mol% CapBioPE, and 0.1 mol% DHPE-Texas Red with Bodipy 505/515-stained TAG (beige) (upper), and a giant unilamellar vesicle (GUV) of the same phospholipid composition (lower). Scale bar is 5 μm. (D) Cryo-electron micrographs depicting an LD with a phospholipid monolayer (upper) and vesicles with a phospholipid bilayer (lower). LDs and vesicles were formed with egg PC. Scale bar is 10 nm.

### Condensation of his-FUS LC on the surfaces of synthetic lipid droplets

We next asked whether lipid droplets (LDs) could act as a surface to catalyze protein phase separation in a similar manner to bilayer surfaces (Fig. 2A). To answer this question, we added N-terminal histidine-tagged proteins to synthetic lipid droplets composed of 83 mol% DOPC, 15 mol% DGS-NTA(Ni), 2 mol% Cap Biotin PE, and 0.1 mol% Texas Red-DHPE and observed their organization upon binding. In our system, DGS-NTA(Ni) was used to recruit different hexahistidine-tagged purified proteins to lipid surfaces to study protein phase separation. To test our ability to recruit his-tagged proteins to the surface of our LDs, we added his-tagged GFP (his-GFP), a protein that does not undergo LLPS, to the LDs. His-GFP covered the surface of the LDs homogeneously (Fig. 2B). Next, we asked how model phase-separating proteins might behave on the LD surface. FUS LC is the low complexity domain of FUS, a well-characterized phase separating protein region that is rich in glutamine, serine, glycine, and tyrosine amino acids. To visualize protein binding, his-tagged FUS LC (his-FUS LC) was labeled with an NHS-ester-reactive ATTO 488 dye on the N-terminus, and samples were imaged under multi-channel confocal microscopy. At a his-FUS LC concentration of 1 μM, a uniform distribution of protein was observed across the surface of the LD (Fig. 2C). When the concentration of his-FUS LC was raised to 2 μM, we observed segregation of the protein into bright and dim regions on the surfaces of LDs (Fig. 2D). Maximum intensity projections of confocal image stacks showed that FUS LC forms round ‘caps’ that are enriched in protein and are surrounded by regions of significantly lower protein intensity (Fig. 2D, rightmost image). The difference between homogeneously bound his-GFP and the binding pattern of his-FUS LC suggests that the intensity variation of FUS LC on the LD surface was not due to phospholipid heterogeneity but rather protein phase separation. In addition, phase separation of FUS LC on the surfaces of LDs appeared morphologically similar to the phase separation of FUS LC on the surfaces of giant unilamellar vesicles (GUVs) of a similar composition (Fig. 2E), which was previously observed (4), suggesting that protein phase separation can occur on both lipid bilayers and monolayers.

**Figure 2:**
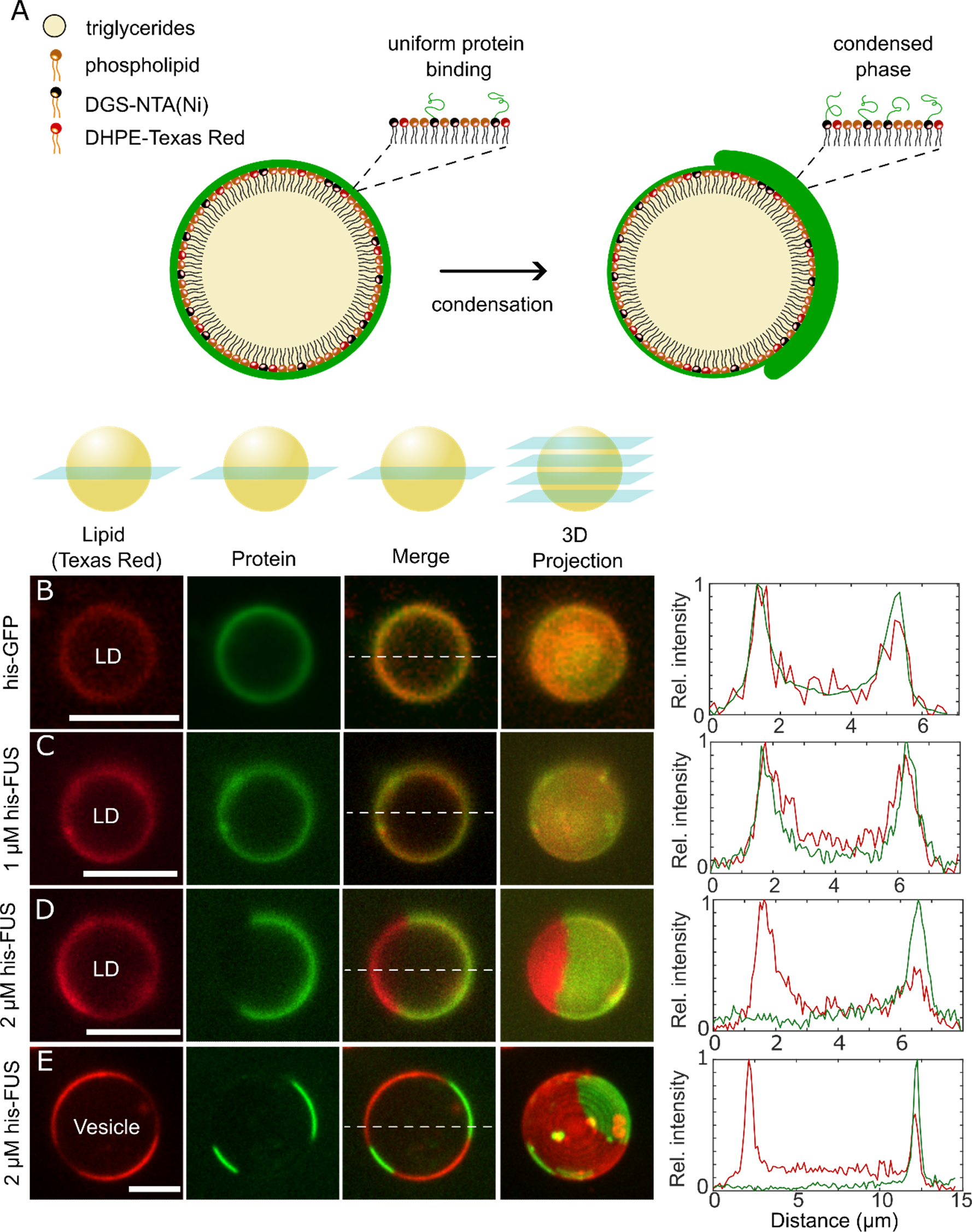
Condensation of FUS LC on the surfaces of synthetic lipid droplets (LDs). (A) Pictorial representation of a uniform protein coat (left) and a condensed protein coat (right) on the LD phospholipid monolayer. Black lipid headgroups represent DGS-NTA(Ni). Red headgroups represent DHPE-Texas Red. Green lines represent his-tagged proteins. (B-E) Representative confocal images (lipid, protein, and merged channels) and corresponding maximum intensity Z-projections of LDs (B-D) or GUVs (E) incubated with the indicated protein in 25 mM HEPES 150 mM NaCl buffer, pH 7.4. The phospholipid composition was 83 mol% DOPC, 15 mol% DGS-NTA(Ni), 2 mol% CapBioPE, and 0.1 mol% DHPE-Texas Red. (B) LDs were incubated with 2 μM his-GFP, which binds uniformly to the LD surface. (C) LDs were incubated with 1 μM his-FUS LC, which covers the LD surface uniformly. (D, E) LDs (D) or GUVs (E) were incubated with 2 μM Atto488-labeled FUS LC, which exhibits protein phase separation on the phospholipid surface. Line profiles depict relative fluorescence intensity of the protein and lipid channels at the locations indicated by the corresponding white dashed lines in (B-E), normalized to the maximum intensity values and then stretched between the maximum and minimum intensity values. Red lines indicate the intensity of the lipid channel and green lines indicate the intensity of the protein channel.

### His-FUS LC condensates on the surfaces of LDs are liquid-like

The boundaries between the protein-poor and protein-rich phases observed in Figure 2 appear circular, which suggests that the shape of the protein phase is controlled by surface tension, a hallmark of liquids (21, 22). To further investigate whether the protein-rich phase formed a liquid-like state, we probed whether FUS LC domains demonstrated additional properties characteristic of liquids, such as merging, re-rounding, and rapid molecular exchange. To observe whether FUS LC domains underwent merging on the LD surface, we imaged LDs immediately after adding protein to the LDs, before protein binding had time to equilibrate. We observed an initially uniform distribution of protein that, within seconds, developed into distinct, rounded, protein-poor regions (Fig. 3A, dark regions) surrounded by a protein-enriched continuous phase. Once these domains grew large enough to be optically resolved, we observed their rapid movement across the LD surface. When two protein-poor domains came into contact, we observed their fusion, followed by re-rounding of the newly formed, larger protein-poor domain within tens of seconds (Fig. 3B) - another hallmark of liquids.

**Figure 3:**
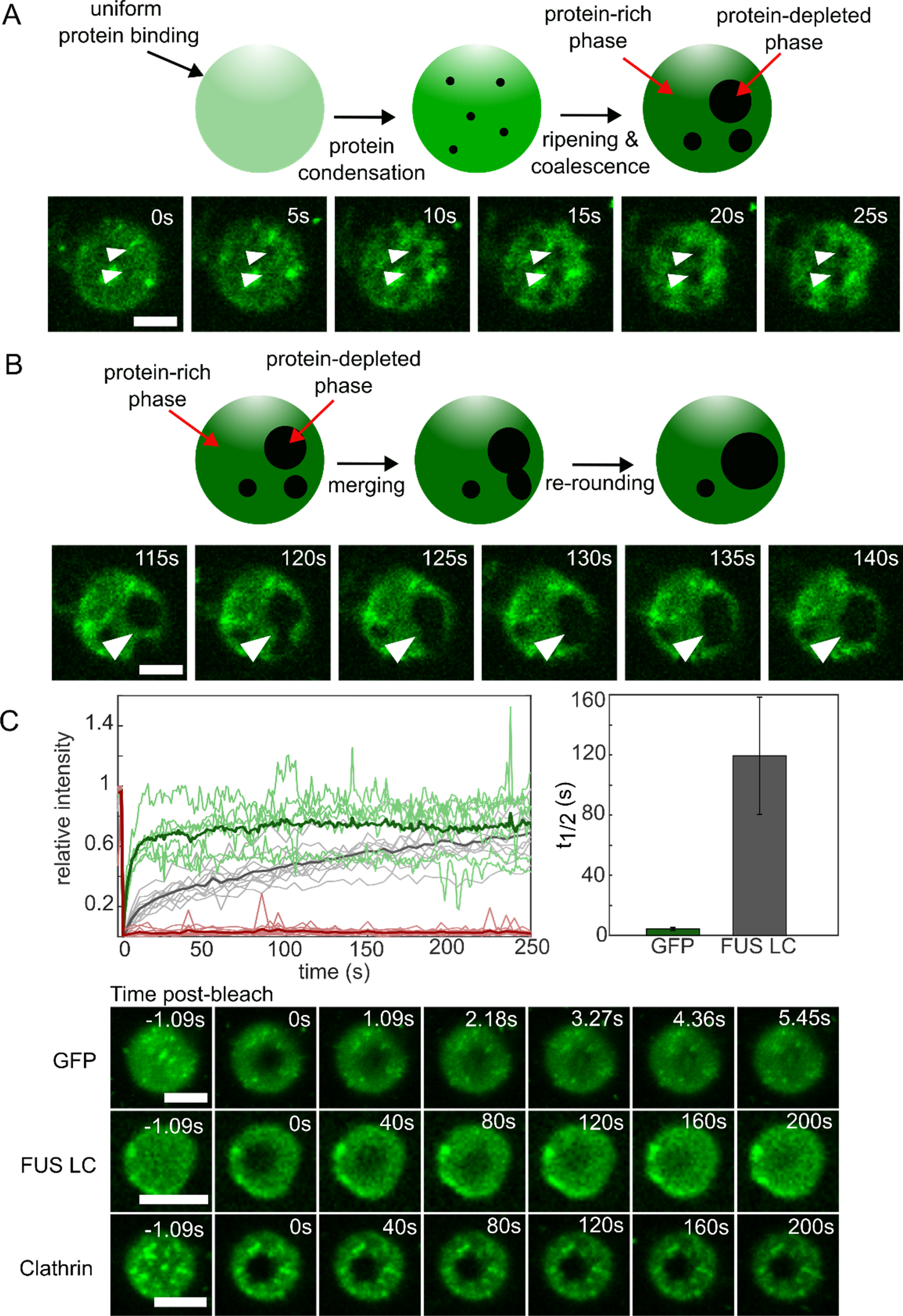
Condensates of FUS LC on the surfaces of LDs are liquid-like. (A) Top: Pictorial representation of the time-dependent emergence of resolvable protein-depleted regions surrounded by a protein-rich continuous phase. Bottom: Confocal image series of the emergence of protein depleted regions on the surface of a LD. White arrows indicate representative regions where protein depletion occurs. Scale bar is 5 μm. (B) Top: Pictorial representation of merging and re-rounding of protein-depleted regions on the LD surface. Bottom: Confocal image series of protein depleted regions merging and re-rounding over time. Scale bar is 5 μm. (C) Top left: FRAP recovery profile after photobleaching a 2.1 μm diameter ROI on lipid droplets. Curves in green represent his-GFP. Curves in gray represent his-FUS LC. Curves in red represent his-Clathrin. Curves in bold indicate the average recovery curve for the corresponding data sets. Top right: t-_1/2_ extracted from the recovery curves for his-GFP (4.28 ± 1.01 s) and his-FUS LC (119.48 ± 39.03 s). T_1/2_ was not calculated for his-Clathrin because essentially no recovery was observed. Data represents mean ± SD (n = 7 LDs for GFP, n = 9 LDs for his-FUS LC, outliers were excluded). Bottom: Image series of fluorescence recovery of LDs covered with his-GFP, his-FUS LC, and his-Clathrin protein. Experiments were conducted in 25 mM HEPES, 150 mM NaCl buffer, pH 7.4. Scale bars are 5 μm.

In addition to merging and re-rounding, liquids often exhibit rapid molecular exchange. We probed the molecular mobility of FUS LC domains on LD surfaces using fluorescence recovery after photobleaching (FRAP). Liquid-like assemblies are formed by a network of weak, multivalent interactions. Therefore, we expected that a liquid-like protein assembly would recover more quickly than a solid-like protein assembly, but more slowly than unassembled proteins that have minimal intermolecular interactions. To test whether protein-rich domains consisting of FUS LC exhibited this intermediate recovery timescale, we compared the recovery profiles of his-FUS LC with his-clathrin and his-GFP. Clathrin, an endocytic coat protein, polymerizes to form a solid-like polyhedral lattice cage (23). In contrast, GFP, which contained the A206K mutation to prevent dimerization (24–26), is a well-structured protein that should diffuse across the LD surface with minimal self-interactions (27). As expected, GFP recovered rapidly after photobleaching (Fig. 3C), with a t_1/2_ of 4.3 ± 1.0 seconds (Fig. 3D), while clathrin showed essentially no recovery (Fig. 3C,E). In contrast, the FUS LC domain exhibited an intermediate recovery rate, with a t_1/2_ of 120 ± 39 seconds (Fig. 3C, D).

### LLPS of his-FUS LC on LDs is driven by increasing salt and protein concentrations

We next investigated whether LLPS on the surfaces of LDs behaves similarly to LLPS of FUS LC in solution (28). Specifically, we mapped the propensity for phase separation on LDs as a function of protein and salt concentration. FUS LC is rich in the amino acids glutamine, glycine, serine, and tyrosine (QGSY), and condensation is driven primarily by hydrogen bonding, π-sp^2^ stacking, π-π, and hydrophobic interactions (28, 29). As the ionic strength of the solution increases, we expect hydrophobic interactions to become more dominant, driving LLPS of FUS LC on the surfaces of LDs (Fig. 4A). All quantifications were made for images collected after 30 minutes of incubation of LDs with protein, followed by 12-15 minutes of imaging. For each salt and protein condition, we observed and categorized three phenotypes of protein binding to LDs: uniform binding, segregation of protein into multiple small domains, and full phase separation into a single, larger protein-rich domain (Fig. 4B). We observed that, as both protein and NaCl concentration increased, the number of LDs with resolvable phase separation increased, in line with previously observed trends (28) (Fig. 4C).

**Figure 4:**
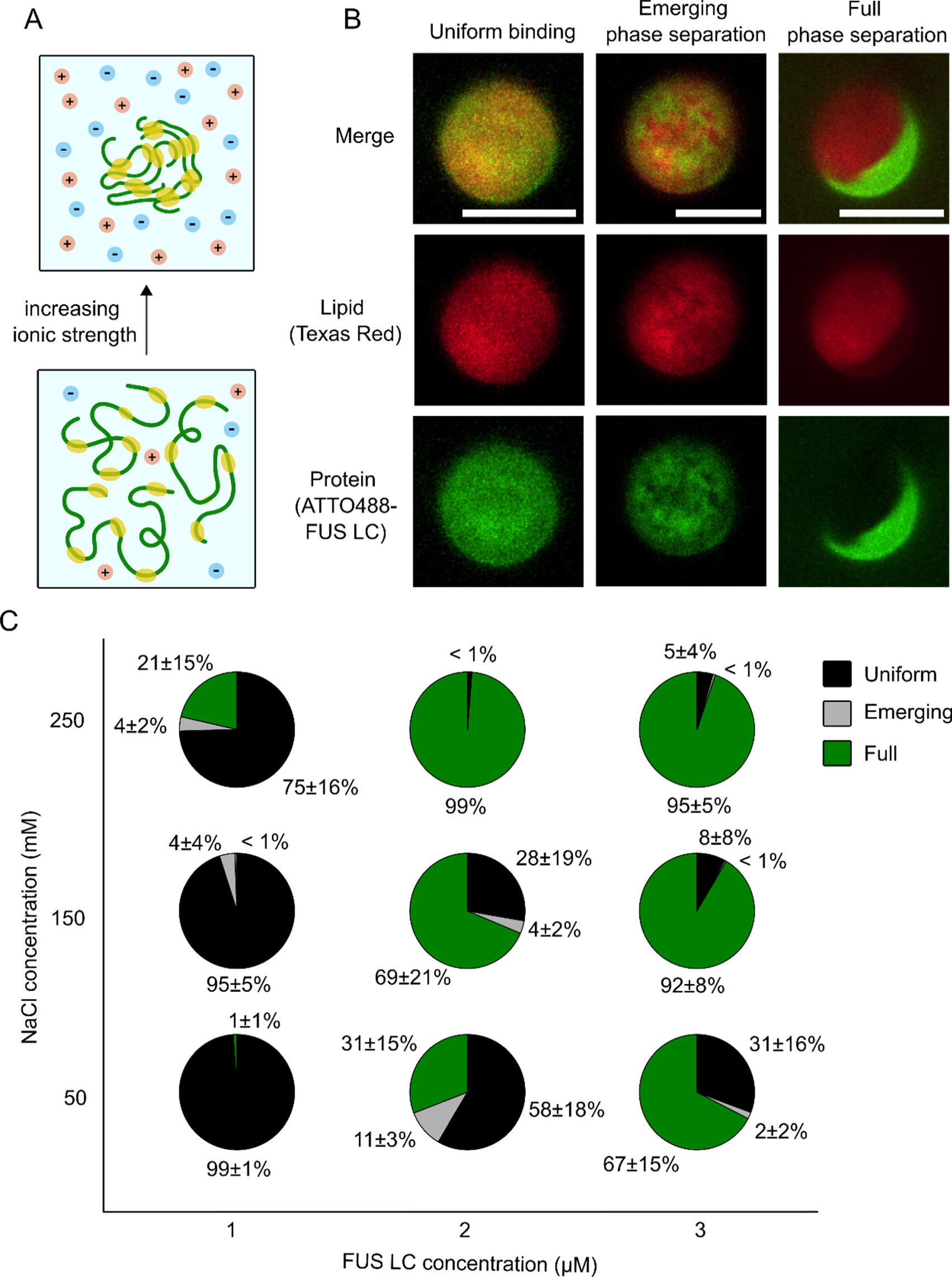
LLPS of his-FUS LC LDs is driven by increasing salt and protein concentrations. (A) Schematic depicting the predicted tendency of his-FUS LC to phase separate as ionic strength of the solution increases. Red and blue dots represent the positively and negatively charged Na+ and Cl-ions, respectively. Green lines represent his-FUS LC chains. Yellow bubbles represent regions of FUS LC self-interaction, such as π-π stacking and hydrophobic interactions. (B) Representative confocal maximum-intensity projections of LDs demonstrating uniform binding (no phase separation, left column), emerging phase separation (center column), and full phase separation (right column). Red channel depicts the lipid channel (DHPE-Texas Red) and the green channel depicts the his-FUS LC protein channel (ATTO-488). Emerging phase separation domains were defined by 4 or more individual resolvable protein-depleted or protein-rich domains. (C) Phase separation diagram mapping the percentage of LDs with uniform protein binding (black), emerging phase separation (gray), and full phase separation (green) under protein concentrations ranging from 1-3 μM and NaCl concentrations ranging from 50-250 mM in 25 mM HEPES, pH 7.4. Data represent the mean ± SEM from >330 LDs from 15 or more fields of view per condition in 3 independent experiments.

### LLPS of his-RGG is screened by increasing ionic strength

To determine whether the ability to undergo LLPS on LD surfaces was unique to FUS LC or could be more universally applicable to other phase-separating proteins, we next tested an additional, well-studied model LLPS protein, which has distinct physicochemical properties relative to FUS LC. Specifically, we studied the RGG domain of LAF1, a DDX3 RNA helicase found in the P granules of *Caenorhabditis elegans* (30). LAF1-RGG is an arginine/glycine rich disordered protein that undergoes electrostatically driven LLPS in solution (30). Indeed, when we incubated his-tagged RGG (his-RGG) with LDs containing DGS-NTA(Ni), we observed LLPS that appeared morphologically similar to that of his-FUS LC (Fig. 5A). When we mapped the phase behavior of his-RGG on the LD surface as a function of ionic strength, we observed a decrease in the number of LDs with LLPS on their surfaces as NaCl concentration increased (Fig. 5B,C). This observation is opposite of the trend that we observed with FUS LC, owing to the difference in the amino acid content of the RGG domain. Specifically, RGG is rich in arginine and glycine residues, so a combination of electrostatic and hydrophobic interactions, such as charge-charge, cation-π, dipole-dipole, and π-π stacking, mediate protein condensation. As salt concentration increases, electrostatic forces weaken due to charge screening, and the hydrophobic effect is enhanced (31). Thus, condensation that is dominated by electrostatic interactions, such as for RGG, is less favorable at higher salt concentrations, while condensation that is dominated by hydrophobic interactions, as is the case for FUS LC, becomes more favorable as ionic strength increases. The alignment of our results with previously observed trends for FUS LC (28) and RGG (32) demonstrates that LLPS on the LD surface is likely driven by the same intermolecular interactions that govern the formation of the respective biomolecular condensates in solution. Collectively, these findings suggest that, similar to bilayer membranes, synthetic LDs can serve as nucleating surfaces for two-dimensional protein phase separation.

**Figure 5:**
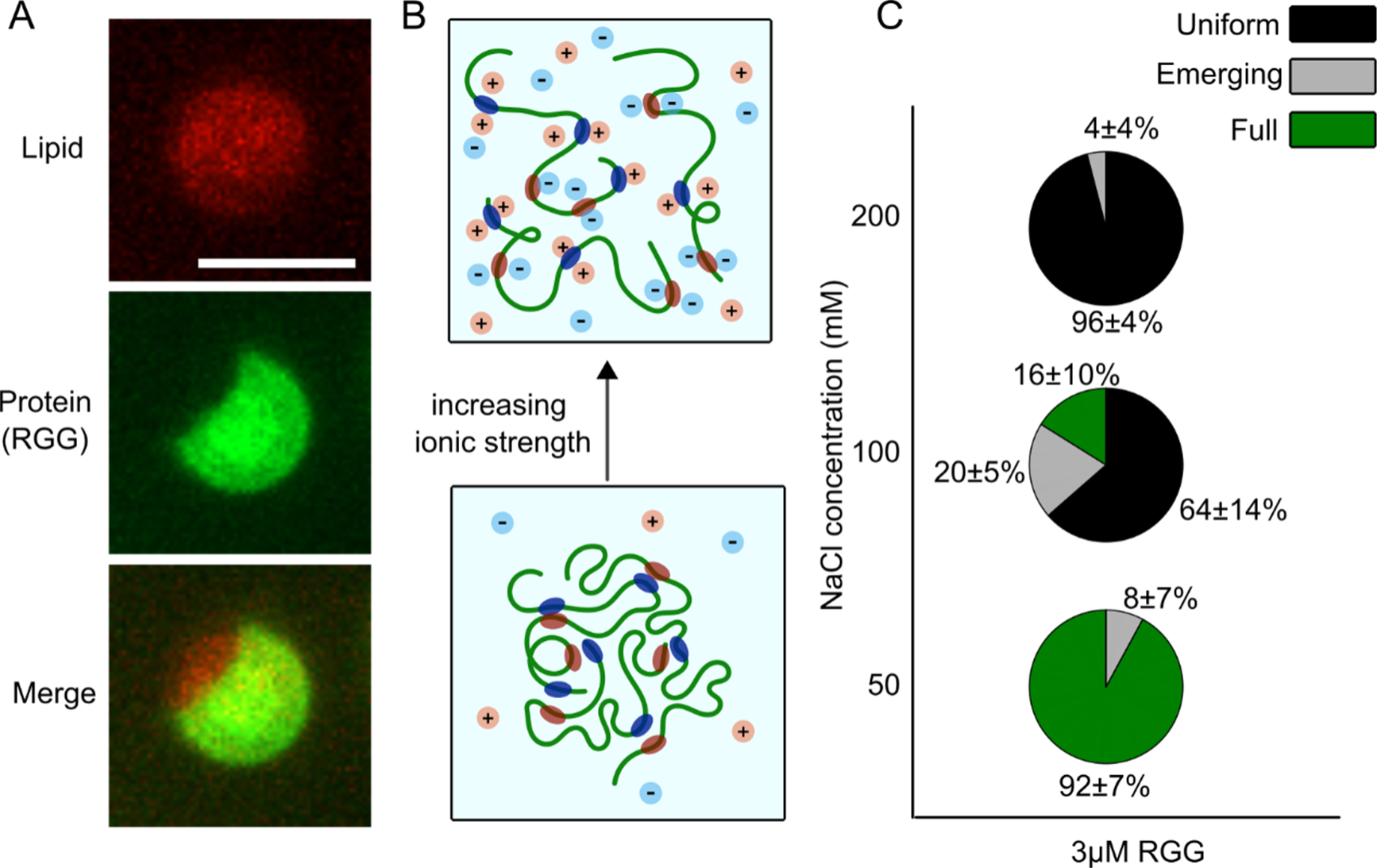
LLPS of his-RGG is screened by increasing ionic strength. (A) Representative maximum intensity projections of ATTO-488 labeled his-RGG phase separation on DGS-NTA(Ni)-containing LDs. Scale bar is 5 μm. (B) Schematic depicting the predicted tendency of his-RGG condensates to dissolve upon increasing ionic strength of the solution. Light red and blue circles in solution represent positively and negatively charged Na+ and Cl-ions in solution, respectively. Green lines represent Atto488-labeled his-RGG protein chains. Dark red and blue ovals attached to his-RGG represent regions of RGG that participate in electrostatic self-interactions. (C) Phase separation diagram mapping the percentage of LDs with uniform binding (black), emerging phase separation (gray), and full phase separation (green) at 3 μM his-RGG and NaCl concentrations ranging from 50-200 mM NaCl in 25 mM HEPES, pH 7.4. Data represent the mean ± SEM from >220 LDs from 15 or more fields of view per condition in 3 independent experiments.

### Condensation of his-FUS LC and his-RGG on cell-derived LDs

The surface of cellular LDs differs from our synthetic LDs, both in lipid composition and in the presence of surface proteins that bind to LDs within cells. For example, perilipin proteins (PLIN1-5), lipases, acyltransferases, and acyl-coA synthetases densely populate the LD surface (33). Therefore, we wondered if protein phase separation was also possible on the surfaces of cellular LDs. To address this question, we isolated LDs from oleate-fed RPE cells (Fig. 6A, panels 1-3) and examined exogenous protein binding to these cell-derived LDs. To enable binding of his-tagged FUS LC and RGG proteins to the cell-derived LD, we incorporated DGS-NTA(Ni) into the LD surface. Before investigating protein binding and condensation, we attempted to visualize whether exogenous lipids could successfully be incorporated onto the cell-derived LD surface through addition of Texas Red-DHPE to cell-derived LDs. When DGS-NTA(Ni) and Texas Red-DHPE were added simultaneously to cell-derived LDs at 14 μM and 1.4 μM concentrations, respectively, a fluorescent ring was clearly observed at the LD surface (Fig. 6A, panel 4). This observation confirms that exogenous lipids were successfully incorporated into the phospholipid monolayer of our cell-derived LDs. We also confirmed binding of his-tagged FUS LC and RGG proteins to the cell-derived LDs, suggesting successful incorporation of DGS-NTA(Ni) (Fig. 6B).

**Figure 6:**
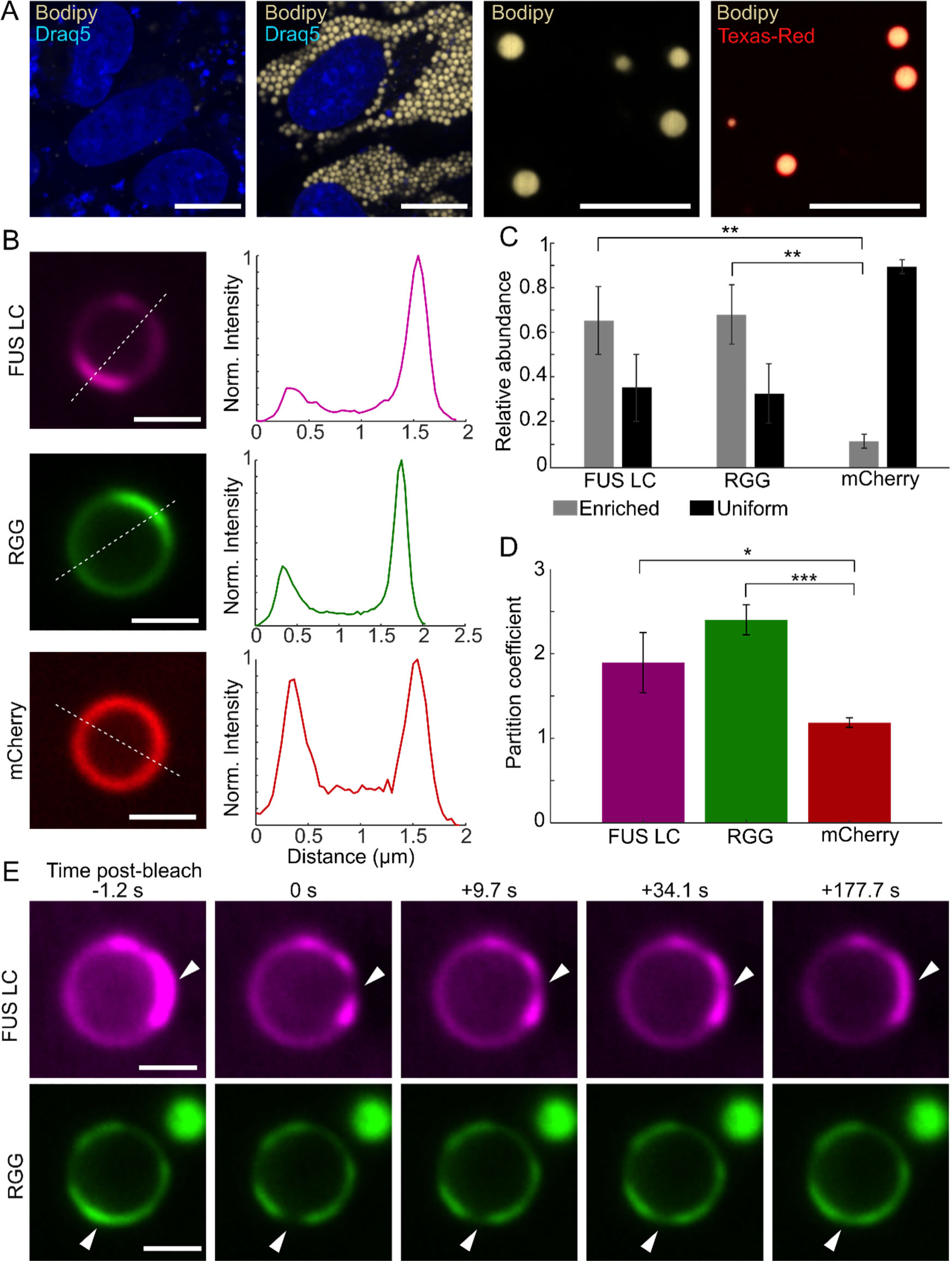
Liquid-liquid phase separation of FUS LC and RGG on cell-derived lipid droplets. (A) From left to right, RPE cells without exogenously added BSA-oleate, RPE cells with 1 mM of exogenously added BSA-oleate, LDs isolated from RPE cells fed with 1 mM exogenous BSA-oleate, and isolated LDs doped with 14 μM DGS-NTA(Ni) and 1.4 μM Texas Red-DHPE. Cells were stained with Draq5 for nuclei (blue) and the neutral lipid dye Bodipy 505/515 for LDs (beige). Cell-derived LDs were stained with Bodipy 505/515. Scale bar for cell images is 10 μm. Scale bar for isolated LD images is 5 μm. (B) Representative images of cell-derived LDs doped with 60 μM DGS-NTA(Ni) in the presence of 5 μM Atto647-FUS LC, Atto488-RGG, or mCherry. Line profiles depict intensity values across the dashed line in the corresponding image. Line profiles were background subtracted and normalized to the minimum and maximum intensity values. Scale bar is 1 μm. (C) Quantification of the relative abundance of uniformly bound and enriched protein for FUS LC, RGG, and mCherry on cell-derived LDs. Data are mean + s.d. across n=3 independent experiments, with at least 28 LDs analyzed per replicate. (D) Average partition coefficient of FUS LC, RGG, and mCherry for the LDs analyzed in C. Partition coefficients were calculated as the ratio of background-subtracted intensity values between the enriched phase and dilute phase, if present, or across two regions of uniform binding. (E) Time series images of fluorescence recovery of FUS LC and RGG enriched regions on the cell-derived LD surface. Timestamps indicate the post-bleach time. Images are representative of recovery behavior observed on at least 3 separate LDs for each protein. Scale bar is 1 μm. Experiments for FUS LC and mCherry on cell-derived LDs were conducted in 25 mM HEPES, 150 mM NaCl, pH 7.4. Experiments for RGG on cell-derived LDs were conducted in 25 mM HEPES, 137 mM NaCl, pH 7.4. For all statistics, one asterisk indicates p<0.05, two asterisks indicate p<0.01, three asterisks indicate p<0.001.

Upon incorporation of DGS-NTA(Ni) and subsequent addition of protein to cell-derived LDs, we observed binding and protein enrichment of FUS LC or RGG, individually, on the cell-derived LD surface, which looked similar to our previous observations of protein phase separation on synthetic LDs (compare Fig. 6B to Fig. 2B). To test whether the enrichment on cell-derived LDs was an effect of uneven DGS-NTA(Ni) incorporation on the LD surface or protein condensation, we compared cell-derived LDs with FUS LC or RGG to cell-derived LDs with his-mCherry (mCherry), a well-structured fluorescent protein that does not undergo protein phase separation. When mCherry bound to our cell-derived LDs, we observed uniform binding to the surface of the majority of LDs, suggesting that the regions of FUS LC and RGG enrichment were an effect of the proteins’ ability to self-interact, rather than a feature of the lipid distribution (Fig. 6B). Notably, protein-enriched regions were observed with around 5-fold greater frequency on the surfaces of LDs exposed to FUS LC or RGG, compared to mCherry, suggesting that these regions were the result of LLPS, rather than non-specific protein aggregation (Student’s t-test, p-values of 0.0037 and 0.0019, respectively) (Fig. 6C). In addition, we calculated the average partition coefficient for each protein on cell-derived LDs. We defined the partition coefficient as the ratio of the maximum intensity to minimum intensity along a line profile through the LD. The line profile was drawn through regions of high and low intensity, if present, or across any location on the LD if binding was visibly uniform. The resulting average partition coefficients for FUS LC and RGG were about 2-fold greater than that of mCherry on cell-derived LDs (Student’s t test, p-values of 0.0270 and 0.0003, respectively) (Fig. 6D). Together, this data strengthened our hypothesis that the regions of FUS LC and RGG enrichment occurred because the proteins self-interact and condense.

We next asked whether regions of FUS LC and RGG enrichment on the surfaces of cell-derived LDs behaved in a liquid-like manner. To assess the mobility of the protein-enriched regions, we bleached small sections of them and observed the fluorescence recovery over time. Within the bleached regions, both FUS LC and RGG visibly recovered within 180 s, suggesting that protein within the enriched phase is mobile (Fig. 6E). Our observations suggest that dynamic protein condensates can form on the surfaces of complex, cell-derived LDs, as well as artificially prepared LDs.

## Discussion

We have demonstrated that protein phase separation can occur on the surfaces of LDs, which are ubiquitous intracellular lipid substrates. Similar to what has been observed previously with lipid bilayers, LLPS occurs on the surfaces of LDs at substantially lower protein concentrations than those required for LLPS in solution. A growing body of recent work has shown that interactions with membranes are an important mechanism by which cells can control LLPS. Specifically, by restricting diffusion of proteins to a two-dimensional surface, membranes can reduce the critical concentration necessary for protein phase separation (34). Here we have shown that the phospholipid monolayer of LDs can perform this same function, which suggests LDs as another candidate lipid substrate for intracellular nucleation of LLPS, in addition to phospholipid membranes. The observed phase separation demonstrates several characteristics of a liquid, including the merging and re-rounding of distinct protein-rich or protein-depleted domains and rapid molecular recovery. This liquid-like behavior suggests that protein condensates on the surfaces of LDs could react quickly to changes in the intracellular environment (34) and organize proteins to catalyze important cellular processes (6).

The LLPS we observed on the surface of LDs was dependent on the protein concentration and ionic strength of the solution. Specifically, when we increased the NaCl concentration from 50 mM to 250 mM, we observed a sharp increase in the number of LDs on which FUS LC underwent LLPS. In contrast, under the same solution conditions, we observed a sharp decrease in the number of LDs on which LAF1-RGG underwent LLPS. Increasing the ionic strength of the solution increases the relative strength of the hydrophobic interactions that drive the condensation of FUS LC, while screening the electrostatic interactions critical for LAF1-RGG condensation. The agreement between these results and the phase behavior previously characterized for FUS LC (28) and LAF1-RGG (32) in solution indicates that proximity to the LD surface does not alter or interfere with the molecular interactions driving phase separation of FUS LC and LAF1-RGG. Further, our observation of FUS LC and RGG LLPS on the surface of LDs isolated from cells shows that, despite the increased complexity of the LD proteome and lipidome compared to reconstituted synthetic LDs, protein LLPS on the lipid monolayer of cell-derived LDs is possible.

It is well-established that LD-associated proteins play important roles in cellular metabolism, including the synthesis and lipolysis of TAGs and protection against cellular stress (35). However, our knowledge and characterization of the structure of many LD-associated proteins remains largely unexplored (36), and many fundamental processes in LD biology, including the process of LD biogenesis and mechanisms of protein targeting to LDs remain unclear (37). For example, LD formation begins with triacylglycerol synthesis by the enzymes DGAT1 and DGAT2 and accumulation of TAGs between the leaflets of the ER membrane to form an oil lens (38). Whether cells have specific sites for lens nucleation and how these sites are determined is still unclear. Similarly, whether and how proteins are involved in determining the direction of LD budding into the cytosol and the scission of LDs from the ER remains to be determined (37).

Could the LLPS of lipid-droplet binding proteins with intrinsically disordered regions (IDRs) play a role in these unanswered questions in LD biology? Building on our results, examining the intersection between the fields of LD cell biology and biophysics could yield candidate proteins for this phenomenon. Indeed, the native LD-binding protein Cidec was recently found to undergo condensation at lipid droplet contact sites despite the apparent lack of an IDR, and its ability to phase-separate was found to be essential for Cidec to perform its function of promoting LD fusion (39). However, Cidec condensation was distinctly gel-like and condensation on other regions of the LD surface was not observed, even when Cidec was bound to the LD at sufficient concentrations (39). Further screening for LD-binding proteins with IDRs could yield proteins that behave as we have observed here with model phase-separating proteins, namely a liquid-like condensation that can spontaneously occur on free LDs, even in the absence of molecular crowders. As a further suggestion that LLPS could play a role in LD biology, local membrane curvature and protein clustering, both of which can be induced by LLPS on membranes (4, 13, 40), are also hypothesized to play a role in organizing the early assembly of LDs (37). Whether LLPS plays a role in these processes remains to be determined. As exploration of the LD proteome continues, the ability of LDs to nucleate phase separation could shed light on multiple facets of LD biology.

## Materials and Methods

### Protein Purification

Expression and purification of his-FUS LC was carried out exactly as described by Yuan et al (4). Briefly, his-FUS LC was overexpressed in E. Coli BL21(DE3) cells. 1-liter cultures were induced with 1 mM IPTG for 4 hours at 37°C and 220 RPM and pellets of cells expressing his-FUS LC were harvested via centrifugation when the OD 600 reached 0.8. The pellets were then lysed in a buffer containing 0.5 M Tris-HCl pH 8, 5 mM EDTA, 5% glycerol, 10 mM β-ME, 1 mM PMSF, 1% Triton X-100 and one EDTA-free protease inhibitor tablet (Sigma Aldrich) for 5 min on ice and then sonicated. The cell lysates were centrifuged at 134,000 x g for 40 min, after which his-FUS LC resided in the insoluble fraction. Therefore, the insoluble fraction was resuspended in a solubilizing buffer of 8 M urea, 20 mM NaPi pH 7.4, 300 mM NaCl and 10 mM imidazole and centrifuged at 134,000 x g for 40 min. In denaturing conditions, his-FUS LC is urea-soluble and thus resided in the supernatant. This supernatant was then mixed with Ni-NTA resin (G Biosciences, USA) for 1 hour at 4°C, settled in a glass column and washed with the above solubilizing buffer. The bound proteins were eluted from the Ni-NTA resin with a buffer containing 8 M urea, 20 mM NaPi pH 7.4, 300 mM NaCl and 500 mM imidazole. In this preparation, the TEV-cleavable his-tag of this protein was not removed, as it was needed for membrane binding via interaction with DGS-NTA(Ni) lipids. The purified proteins were then buffer-exchanged into 20 mM CAPS pH 11 storage buffer using 3K Amicon Ultra centrifugal filters (Millipore, USA). Small aliquots of the protein were frozen in liquid nitrogen at a protein concentration of approximately 1 mM.

Expression and purification of his-LAF-1 RGG was performed exactly as described previously (4). Briefly, his-LAF-1 RGG was overexpressed in E. Coli BL21(DE3) cells. 1-liter cultures were induced with 0.5 mM IPTG overnight at 18°C and 220 RPM, and pellets of cells expressing his-LAF1 RGG were harvested from the cultures via centrifugation when the OD 600 reached 0.8. The pellets were then lysed in a buffer containing 20 mM Tris, 500 mM NaCl, 20 mM imidazole, and one EDTA-free protease inhibitor tablet (Sigma Aldrich) for 5 min on ice and then sonicated. The cell lysates were centrifuged at 134,000 x g for 40 min, after which His-LAF-1 RGG resided in the supernatant. The supernatant was then mixed with Ni-NTA resin (G Biosciences, USA) for 1 hour, settled in a glass column, and washed with a buffer containing 20 mM Tris, 500 mM NaCl, 20 mM imidazole, at pH 7.5. The bound proteins were eluted from the Ni-NTA resin with a buffer containing 20 mM Tris, 500 mM NaCl, 500 mM imidazole, at pH 7.5. The purified proteins were then buffer-exchanged into 20 mM Tris, 500 mM NaCl, at pH 7.5 using Amicon Ultra centrifugal filters. Small aliquots of the protein were frozen in liquid nitrogen at a protein concentration of approximately 120 μM and stored at −80 °C. To promote solubility of the LAF-1-RGG protein, the entire purification process was performed at room temperature, except for the cell lysis process, which was done on ice.

His-clathrin was purified according to a previously published protocol (41). His-GFP and his-mCherry purification were performed according to a previously published protocol (42).

### Protein Labeling

His-FUS LC was labeled with Atto-488 or Atto-647 NHS-ester fluorescent dyes (ATTO-TEC, Sigma-Alrich). Labeling occurred at or near the N-terminus because only the N-terminus and a lysine at residue position 5 in the leader sequence preceding the N-terminal region hexa-histidine tag were expected to react with the NHS-functionalized dye. The labeling reaction took place in a 50 mM HEPES buffer at pH 7.4. Dye was added to the protein in 2-fold stoichiometric excess and allowed to react for 30 min at room temperature, empirically resulting in a labeling ratio near 1:1 dye:protein. Labeled protein was then buffer-exchanged into 20 mM CAPS pH 11 buffer and separated from unconjugated dye using 3K Amicon columns (MilliporeSigma). His-LAF-1 RGG was labeled with Atto-488 in its storage buffer (20 mM Tris, 500 mM NaCl, pH 7.5). Protein and dye concentrations were monitored using UV–Vis spectroscopy using a molar extinction coefficient of 35760 cm^-1^ for FUS LC and 16390 cm^-1^ for RGG. Labeled proteins were dispensed into small aliquots, flash frozen in liquid nitrogen and stored at −80°C. His-Clathrin was labeled with Atto594 NHS-ester (ATTO-TEC, Sigma-Aldrich) according to a previously published protocol (41) and was ultracentrifuged and used immediately after labeling to minimize aggregation.

Labeled protein was mixed with unlabeled protein (if necessary) to achieve a 10% labeled protein stock for use in all imaging experiments.

### Preparation of Synthetic Lipid Droplets

1,2-dioleoyl-sn-glycero-3-phosphocholine (DOPC), 1,2-dioleoyl-sn-glycero-3-phospho-(1’-rac-glycerol) (sodium salt) (DOPG), L-α-phosphatidylcholine (Egg PC), 1,2-dioleoyl-sn-glycero-3-[(N-(5-amino-1-carboxypentyl)iminodiacetic acid)succinyl] (nickel salt) (18:1 DGS-NTA(Ni)), and 1,2-dioleoyl-sn-glycero-3-phosphoethanolamine-N-(cap biotinyl) (sodium salt) (18:1 Biotinyl Cap PE) were purchased from Avanti Polar Lipids. Texas Red-DHPE was purchased from AAT Bioquest. All lipid stocks were stored at −80°C.

Glass vials were cleaned by washing with acetone, then with Neutrad (Fisher Scientific), and rinsing with ultrapure H_2_O. Lipid mixtures dissolved in chloroform were mixed in the appropriate molar ratios in the clean glass vial to obtain 1.60×10^-7^ moles of lipid. 10 μL of olive oil was added to the lipid mixture and gently shaken to mix before the chloroform was evaporated under a stream of N_2_. The lipids were further dried in a vacuum desiccator for at least 2 hours to remove all solvent. Lipids and olive oil were then heated at 37°C for 10 minutes and vortexed for 10 seconds to evenly mix the lipids within the olive oil. The lipid mixture was resuspended in 25mM HEPES, 150mM NaCl (pH 7.4) and vortexed 3 times for 10 seconds to generate an emulsion of lipid droplets suspended in aqueous buffer. Lipid droplets were stored at 4°C and used within one day of formation.

### Preparation of GUVs

GUVs were prepared as previously reported using electroformation (4). Lipid mixtures dissolved in chloroform were combined in a glass vial washed with acetone and Neutrad and rinsed with ultrapure H_2_O to obtain 2.0×10^-7^ moles of lipid. The lipid mixture was spread into a film on acetone- and Neutrad-cleaned indium-tin-oxide (ITO) coated glass slides (resistance ∼8-12W sq^-1^) and further dried in a vacuum desiccator for at least 2 hours to remove all the solvent. Electroformation was performed at 55°C in 340 milliosmole glucose solution. The voltage was increased every 3 min from 50 to 1400 mVpp for the first 30 min at a frequency of 10 Hz. The voltage was then held at 1400 mVpp for 120 min and finally was increased to 2200 mVpp for the last 30 min during which the frequency was adjusted to 5 Hz. GUVs were stored at 4°C and used within 2 days after electroformation.

### Tethering of Synthetic Lipid Droplets and GUVs

LDs and GUVs were tethered to glass coverslips as previously described (4). Briefly, glass coverslips were passivated with a layer of 2% biotinylated PLL-PEG8kDa. LDs and GUVs were doped with 2% 18:1 biotinyl Cap PE (Avanti Polar Lipids) and tethered to the passivated surface using NeutrAvidin. PLL-PEG was synthesized by combining amine reactive PEG and PEG-biotin in molar ratios of 98% and 2%, respectively. This PEG mixture was added to a 20 mg/mL mixture of PLL in 50 mM sodium tetraborate (pH 8.5) such that the molar ratio of lysine subunits to PEG was 5:1. The mixture was continuously stirred at room temperature overnight then buffer exchanged into 25 mM HEPES, 150 mM NaCl (pH 7.4) using Centri-Spin size exclusion columns (Princeton Separations).

To each 5 mm imaging well, 20 μL of 2% biotinylated PLL-PEG was added and incubated at room temperature for 20 min. Wells were washed with 25 mM HEPES (pH 7.4) containing the NaCl concentration specified for the experiment using gentle pipetting until PLL-PEG reached at least a 1000-fold dilution. Then 4 μL of NeutrAvidin at a stock concentration of 1μg/μL dissolved in 25mM HEPES 150mM NaCl (pH 7.4) was added to the well, incubated at room temperature for 10 min, and wells were washed with the appropriate buffer to remove excess Neutravidin (1000-fold dilution). LDs or GUVs were diluted in a 1:5 or 1:6 ratio, respectively, added to the well, and incubated for 10-15 minutes at room temperature. For LDs, the glass slide was turned upside down to facilitate adherence to the treated surface as the LDs tended to rise upwards within the aqueous buffer. For GUVs, the well was gently rinsed with the appropriate buffer to remove excess GUVs. For experiments involving protein, protein was then added to the well at the appropriate concentration, and wells were covered with a glass coverslip to prevent evaporation prior to imaging using confocal microscopy.

### Fluorescence Microscopy

Imaging experiments were performed using a spinning disc confocal super resolution microscope (SpinSR10, Olympus, USA) equipped with a 1.49 NA/100X oil immersion objective fitted with a Hamamatsu Orca Flash 4.0V3 scientific complementary metal–oxide–semiconductor digital camera. Laser wavelengths of 488, 561, and 640 nm were used for excitation. For sample preparation, No. 1.5 Glass coverslips were cleaned in 2% v/v Hellmanex III (Hellma Analytics) solution, rinsed thoroughly with ultrapure H_2_O, and dried with compressed air. Imaging wells were formed by cutting 5 mm holes in PDMS gaskets or 1.5-mm thick silicone gaskets (Grace Biolabs) using a biopsy punch. Gaskets were cleaned with Neutrad, rinsed with ultrapure H_2_O, and dried thoroughly with compressed air before being placed on dry coverslips to create a temporary waterproof seal. A second coverslip was placed on top of the imaging well to seal the chamber and prevent evaporation.

### Cryo-Electron Microscopy

To image the synthetic LDs using cryo-EM, a diameter of 50-200 nm and a LD high concentration was desired. To achieve size reduction and increased concentration of LDs, 2.46 mg of egg PC and 14 µL of olive oil were dried overnight in a vacuum desiccator. Lipids and olive oil were rehydrated in 25 mM HEPES 150 mM NaCl, pH 7.4 by vortexing and tip sonicated on ice for 5 cycles of 1 minute on ice. 2.5 µl of LDs was applied to glow discharged holey carbon grids (Quantifoil 1.2/1.3), blotted for 6 s with a blot force of 0 and rapidly plunged into liquid nitrogen-cooled ethane using an FEI Vitrobot MarkIV. Images were acquired on a FEI Glacios cryo-TEM equipped with a Falcon 4 detector with a pixel size of 0.94 Å, and a total exposure time of 15s resulting in a total accumulated dose of 40 e/Å^2^ which was split into 60 EER fractions. Motion correction was performed using crypSPARC(43).

For the quantification of monolayer thickness, the thickness of the dark phospholipid ring was measured using ImageJ. 17 LDs were measured at three separate locations for each LD structure for a total of 51 measurements. All 51 measurements were averaged, as shown in Figure S1C. For the quantification of bilayer thickness, the total distance across two resolvable phospholipid layers was measured using ImageJ. 20 vesicles were measured at three separate locations for each vesicle structure, for a total of 60 measurements. All 60 measurements were averaged, as shown in Figure S1C.

### Phase Diagram Experiments

For phase diagram experiments involving varying NaCl concentrations, wells were washed, and protein dilutions were made with 25 mM HEPES with a NaCl concentration corresponding to the desired final concentration. For his-FUS LC experiments, initial protein dilutions were made in 0.5 mM CAPS, 150 mM NaCl to prevent droplet formation in solution. Final protein dilutions prior to addition to the well were made in 25 mM HEPES with a NaCl concentration that matched the desired NaCl concentration. For his-RGG experiments, initial protein dilutions were made in 50 mM Tris, 500 mM NaCl (pH 7.4) to prevent droplet formation in solution. Final protein dilutions prior to addition to the well were made in 25 mM HEPES with an NaCl concentration adjusted such that the final concentration of the protein dilution matched the desired NaCl concentration. Following protein addition, samples were sealed to prevent evaporation and allowed to incubate for 30 minutes before imaging. Image stacks taken at fixed distances perpendicular to the coverslip plane were acquired after protein addition. At least 15 fields of views were randomly selected for each sample for further analysis after the addition of protein.

### Fluorescence Recovery After Photobleaching on LDs

For experiments on synthetic LDs, fluorescence recovery after photobleaching was performed on an Olympus Fluoview 3000 confocal microscope with a 1.1 NA/60X water immersion objective and water-matching oil immersion medium (n=1.3310, Cargille Laboratories). Bleaching of a 2.1 μm diameter ROI was performed using a 488 nm laser at 75% laser power for 10 seconds per ROI. Images were acquired every 1.09 s for his-GFP and every 5 s for his-FUSLC and his-Clathrin to capture the dynamics of recovery while minimizing photobleaching. Raw his-GFP and his-FUS LC intensity values for the bleached region were normalized to account for photobleaching using **Equation 1** and stretched to standardize bleach depth using **Equation 2**.

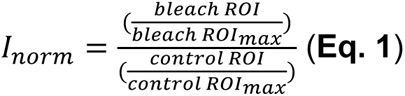

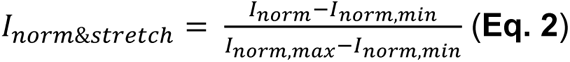

Raw intensity values for the bleached region for his-Clathrin were not normalized for photobleaching because while no recovery was observed, photobleaching of the surrounding region caused an artificial rise in recovery using the previous two formulas. Thus, his-Clathrin raw intensity values were normalized using **Equation 3** and stretched using **Equation 2**.

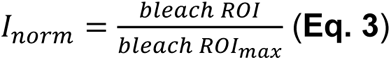

Normalized and stretched recovery curves were plotted as a function of time. Average recovery curves for each protein were calculated by averaging the normalized and stretched intensity values for all replicates at each time step.

#### Calculation of t_1/2_

To calculate t_1/2_ values for his-FUS LC, the data for each replicate was fit to the double-exponential equation of **Equation 4** using the Levenberg-Marquardt method with an initial parameter guess of 1 for each coefficient in MATLAB:

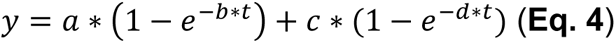

The equation was solved for the time at which the intensity reached 0.5 using the obtained fit coefficients to calculate t_1/2_.

Because the his-GFP data was noisy and the data reached a plateau of recovery within the timespan of data collection, fitting was not needed to calculate the t_1/2_ values. Instead, the times and intensities immediately before and after the intensity reached 50% of the maximum intensity were used to linearly interpolate the time at which the intensity reached 0.5*(max intensity).

For experiments on cell-derived LDs, FRAP was performed using the Olympus FRAP unit 405 nm laser and an additional 3.2X magnification lens using 32% laser power.

### Mammalian cell culture

Human retinal pigmented epithelial (RPE) cells (ARPE-19) were purchased from American Type Culture Collection (Manassas, VA). RPE cells were cultured in 1:1 F12:DMEM supplemented with 10% FBS, 20 mM HEPES, and 1% penicillin, 1% streptomycin, 1% L-glutamine (PSLG). Cells were incubated at 37 °C with 5% CO_2_ and passaged every 72-96 hours. If cells were incubated for 96 hours between passages, media was changed at the 48-hour time-point.

For fluorescent imaging, cells were seeded in chambered 8-well glass bottom slides (CellVis) at a density of 5,000 cells per well. For cells in which LD formation was induced, cell media was changed and a final concentration of 1 mM OA-BSA was added to culture media for 48 hours before imaging. Prior to imaging, the media was removed, and cells were washed with PBS and incubated in PBS containing 5 μM Draq5 far-red nuclear stain (Fisher Scientific) and 2 μM Bodipy 505/515 (Fisher Scientific) or 10 μM Nile Red (Sigma-Aldrich) neutral lipid stain for 10 minutes. Cells were washed again with PBS and fresh phenol red-free media was added prior to imaging.

### Isolation of cell-derived lipid droplets

To induce LD formation in RPE cells, oleic acid (OA) (Sigma-Aldrich) was complexed to fatty acid-free bovine serum albumin (BSA) (Sigma-Aldrich) at a 5:1 molar ratio of OA:BSA to form OA-BSA. To prepare a 15 mM stock of OA-BSA, first a 300 mM stock of OA was prepared by dissolving oleic acid in EtOH in a glass scintillation vial. When not in use, OA and stocks were flushed with N_2_ and kept at −20°C. The appropriate mass of BSA was weighed out and dissolved in cell culture media (1:1 F12:DMEM + 10% FBS) warmed to 37°C. The BSA solution was mixed on a stir plate at 37°C for 30 minutes to ensure dissolution of BSA in the media. After BSA dissolved in warm media, additional media was added to achieve the desired stock concentration of 3 mM BSA, and 300 mM OA was added to the BSA solution for a final OA concentration of 15 mM, yielding a 5:1 ratio of OA:BSA. OA precipitates upon addition to the BSA solution, and dissolution was achieved through stirring overnight at 37°C at 400 rpm on a stir plate. After OA was dissolved in BSA-containing media, the OA-BSA solution was sterile filtered, aliquoted, and stored at −20°C until use. OA-BSA aliquots were thawed at room temperature and used only once to avoid freeze-thaw cycles. Stock concentrations were calculated such that the final EtOH concentration was 0.5% v/v when OA-BSA was added to cells at a final concentration of 1 mM OA.

To induce LD formation in RPE cells, RPE cells were passaged and seeded into a 150 mm round cell culture flask. After 48 hours, the culture media was exchanged and 15 mM OA-BSA was added to a final concentration of 1 mM OA. Cells were incubated for an additional 48 hours, after which time cells were ∼90% confluent and LDs were densely packed in the cytoplasm.

LDs were isolated according to a previously published STAR protocol (44), with some modifications. Briefly, cells were gently washed with PBS and collected in PBS using a cell scraper. Cells were centrifuged at 500 x g for 10 min at 4°C to pellet cells, and the supernatant was discarded. The cell pellet was resuspended in 2 mL of chilled 20 mM Tris 1 mM EDTA (referred to as HLM buffer) containing 1X Pierce protease inhibitor (Thermo Fisher Scientific) and incubated on ice for 10 min before homogenization using a Dounce homogenizer. Homogenate was centrifuged at 1000 x g for 10 min at 4°C to pellet nuclei and unbroken cells. The supernatant was collected and ultracentrifuged within a sucrose density gradient to concentrate the LDs and separate them from the ER and other cell compartment fractions. To create the sucrose gradient, the 2 mL of supernatant was thoroughly mixed with 1 mL of 60% sucrose in HLM buffer to create a final concentration of 20% sucrose containing the cell homogenate and was added to a clean and dry ultracentrifuge tube. 2.1 mL of 5% sucrose HLM buffer was layered on top carefully, so as not to disturb the lower layer. 1.89 mL of HLM buffer was then carefully layered on top of the 5% sucrose layer for a total volume of 7 mL. The sample was centrifuged at 28,000 x g in a S50-ST swinging bucket rotor (Thermo Fisher Scientific) in a Sorvall MX 150 Plus Micro-Ultracentrifuge (Thermo Fisher Scientific) for 30 min at 4°C to float LDs. After centrifugation, LDs were concentrated at the top of the tube and were collected by pipette for imaging and used within 72 hours of isolation.

### Cell-derived LD imaging

To image LDs without the exogenous addition of protein, a 2 μM final concentration of Bodipy 505/515 was added to the solution of LDs and gently mixed before adding to the imaging chamber. The imaging chamber was inverted and incubated for 10 minutes to facilitate LD adhesion to the clean coverslip surface before the chamber was covered with an additional coverslip to prevent evaporation and cell-derived LDs were imaged.

To enable His-tagged protein binding to the cell-derived LD surface, cell-derived LDs were mixed with a 2 mM stock concentration of DGS-NTA(Ni) (Avanti Polar Lipids) dissolved in THF for a final concentration of 60 μM DGS-NTA(Ni) and a final v/v concentration of 3% THF. The volume of DGS-NTA(Ni) added was limited to 3% v/v to minimize any possible disruption to the LDs in the presence of an organic solvent. Cell-derived LDs were vortexed 5 times for 5-second intervals to facilitate incorporation of DGS-NTA(Ni) into the lipid monolayer, as previously demonstrated with biotinylated lipids and giant plasma membrane-derived vesicles (45). Ni-NTA-doped cell-derived LDs were added to the imaging chamber and the chamber was inverted and incubated for 10 minutes. Then the chamber was washed gently 3 times with 60 μL of 25 mM HEPES to remove excess THF and unincorporated DGS-NTA(Ni). To facilitate faster diffusion and protein mixing, protein was first diluted with buffer (25 mM HEPES, pH 7.4 at 150 mM NaCl for FUS LC and mCherry or 137 mM NaCl for RGG) outside of the imaging chamber to achieve a larger volume of protein, then added to the imaging chamber to achieve the desired final concentration. Protein was gently mixed upon addition to the chamber and the chamber was sealed with an additional coverslip to prevent evaporation during imaging.

