## Supplemental figures for "Lipid droplets as substrates for protein phase separation"

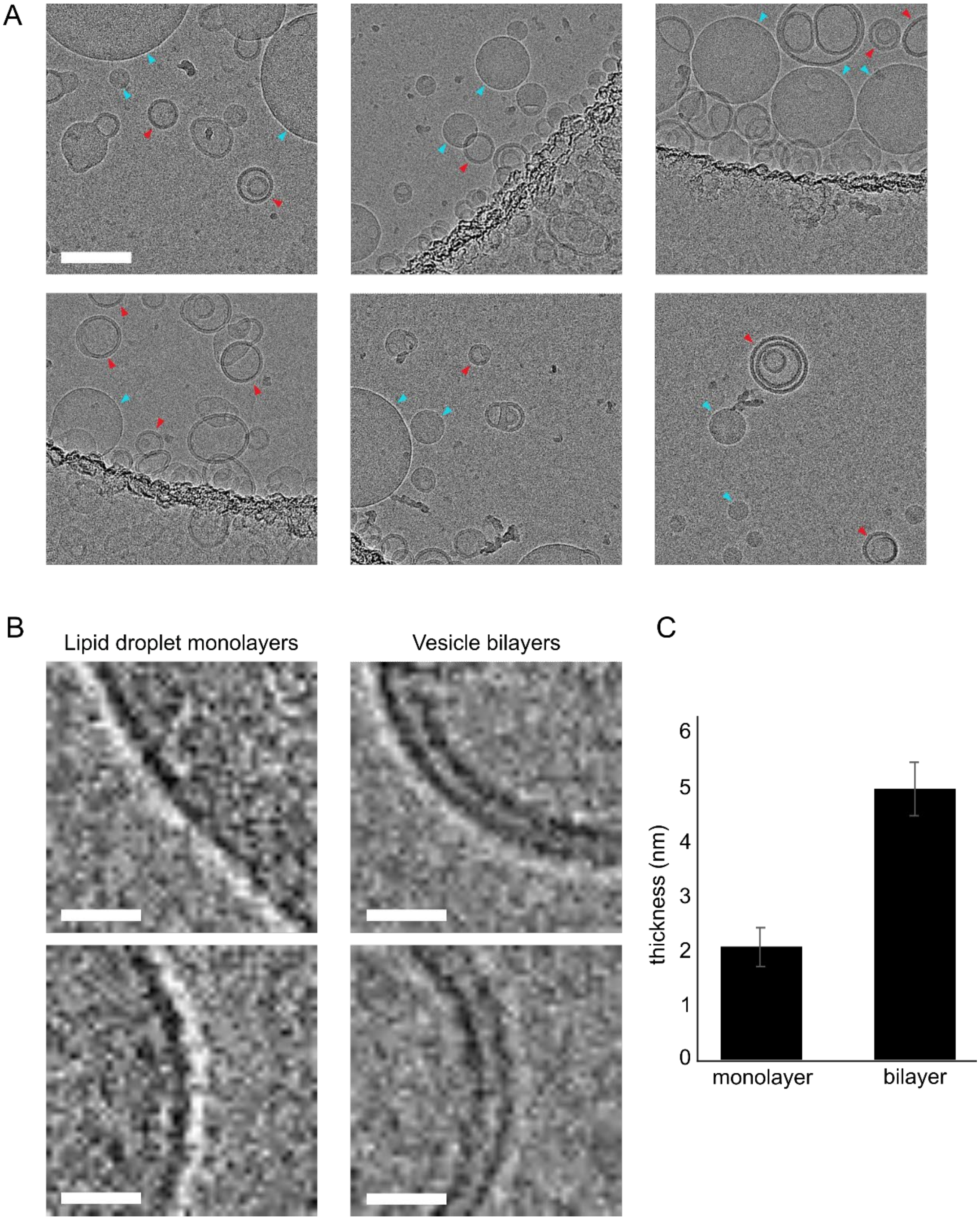


**Figure S1: Cryo-EM characterization of synthetic lipid droplets.** (A) Additional micrographs of samples containing bilayer and multilamellar vesicles and monolayer lipid droplets. Blue arrowheads point to structures identified as lipid droplets with one visible layer of phospholipids. Red arrowheads point to structures identified as vesicles with two visible layers of phospholipids. Scale bar is 100 nm. (B) Zoomed-in images of regions of lipid droplet monolayers showing a single dark line of phospholipids and bilayer vesicles showing two clearly resolvable dark lines of phospholipids. Scale bar is 10 nm. (C) Quantification of the thickness of monolayers and bilayers measured from cryo-EM micrographs. The average monolayer thickness is 2.1 ± 0.3 nm (n = 17 LDs). The average bilayer thickness is 5.0 ± 0.5 nm (n=20 vesicles).

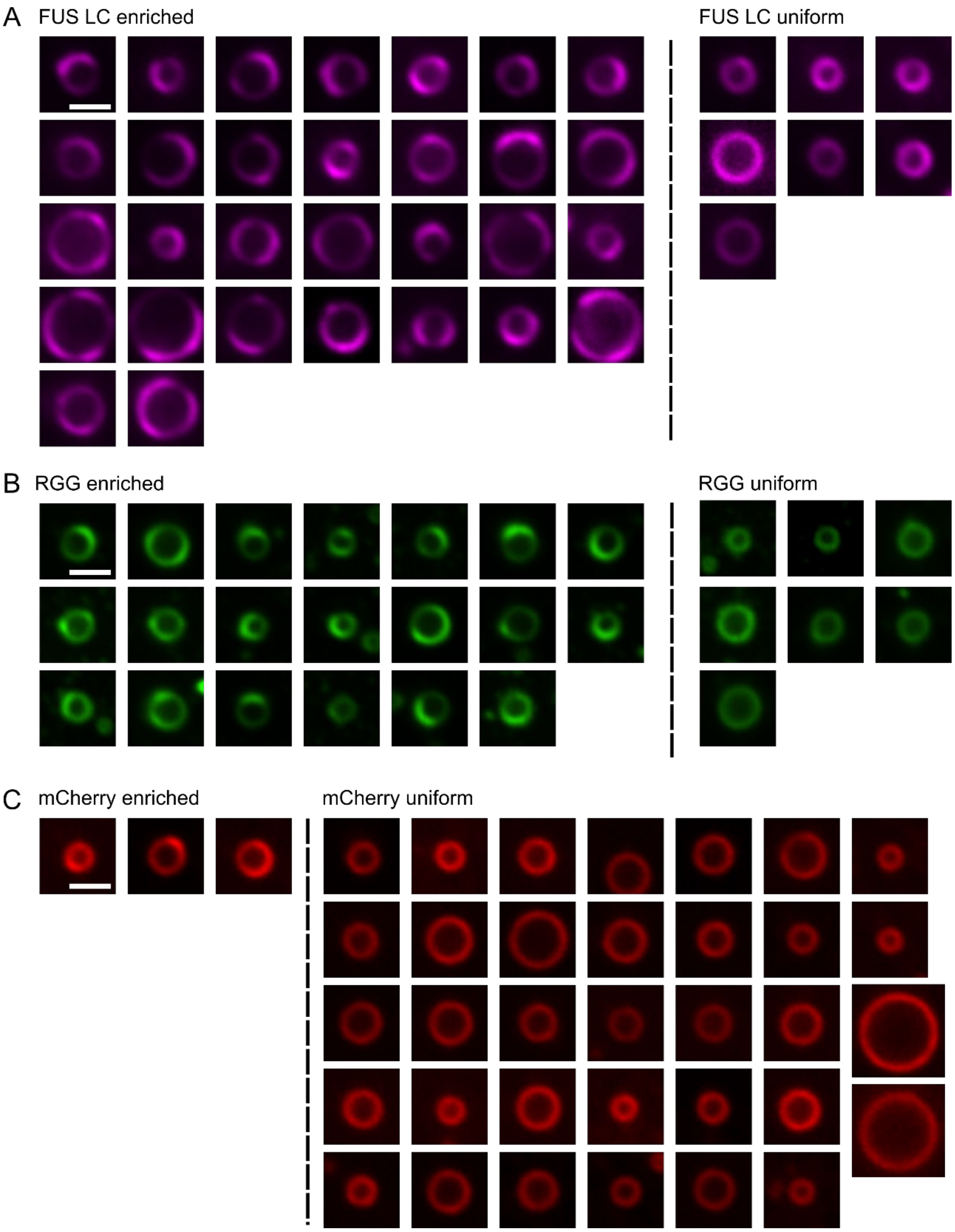


**Figure S2: Cell-derived LDs with his-recruited proteins.** (A) All images that were analyzed for one of three replicates of FUS LC binding to cell-derived LDs. (B) All images that were analyzed for one of three replicates of RGG binding to cell-derived LDs. (C) All images that were analyzed for one of three replicates of mCherry binding to cell-derived LDs. LDs are categorized as enriched or uniform based on their observable patterns of protein binding. All scale bars are 1 μm.
